# Marker-free imaging of α-Synuclein aggregates in a rat model of Parkinson’s disease using Raman microspectroscopy

**DOI:** 10.1101/2021.02.02.429468

**Authors:** Fide Sevgi, Eva M Brauchle, Daniel A Carvajal Berrio, Katja Schenke-Layland, Nicolas Casadei, Madhuri S Salker, Olaf Riess, Yogesh Singh

## Abstract

A hallmark of Parkinson’s disease (PD) is the formation of Lewy bodies in the brain. Lewy bodies are rich in the aggregated form of misfolded α-Synuclein (α-Syn). The brain from PD patients can only be analysed after post-mortem, limiting the diagnosis of PD to the manifestation of motor symptoms. In PD patients and animal models phosphorylated α-Syn was detected in the gut, thus, raising the hypothesizes that early-stage PD could be diagnosed based on colon tissues biopsies. Non-invasive marker-free technologies represent an ideal method to potentially detect aggregated α-Syn *in vivo.* Raman microspectroscopy has been established for the detection of molecular changes such as alterations of protein structures. Here, the olfactory bulb in the brain and the muscularis mucosae of colon tissue sections of a human BAC-SNCA transgenic (TG) rat model was analysed using Raman imaging and microspectroscopy. Raman images from TG and WT rats were investigated using spectral, principal component and true component analysis. Spectral components indicated protein aggregates (spheroidal oligomers) in TG rat brain and colon tissues even at a young age but not in WT. In summary, we have demonstrated that Raman imaging is capable to detect α-Syn aggregates in colon tissues of a PD rat model and making it a promising tool for future use in PD pathology.

## Introduction

Parkinson’s disease (PD) is the second most common disorder among neurodegenerative diseases with 6.1 million persons afflicted worldwide as estimated in 2016 (1, 2). This disease burden is projected to have doubled in the past 25 years, while the number of older people did not increase in the same amount, indicating environmental factors could have an important role in PD progression (2). PD is manifested by the loss of neurons in the *substantia pars nigra compacta* with an increased neural loss of up to 70% by the time of death (3).

The presence of Lewy bodies (LB) represents the pathological hallmark of PD, as they were linked to the death of the dopamine producing cells in the brain (4). The major component of LB is the filamentous inclusion protein α-Synuclein (α-Syn) (5). Accurate process of *in vivo* LB formation is not known. However, it is widely accepted that aggregation of α-Syn into soluble oligomers and then insoluble amyloid fibrils is the foundation of LB (6). During the aggregation process, phosphorylation is a usual characteristic as post-translational phosphorylation of α-Syn is observed in 90% of misfolded proteins, while in cytosolic α-Syn only 4% is phosphorylated (7). Even though α-Syn is an abundant protein in the brain, its exact function remains elusive.

In its natively unfolded nature, α-Syn is a monomeric or intrinsically disordered protein in neuronal cells and highly conserved protein and mostly found in the presynaptic terminals of neurons and possibly in the nucleus (5, 8, 9). However, α-Syn adopts an α-helical nature upon engaging with lipid membranes and detergent micelles (10, 11). Nuclear magnetic resonance (NMR) studies also demonstrated that the N-terminal region of the protein had a tendency towards forming stable α-helical secondary structures (12, 13). Further, *in vitro* studies suggested that monomeric α-Syn was consisting mostly of α-helical (49%) and extended β-strand and polyproline II (PPII) structures (41%) with only a small amount of β-sheet present (10%) (14). Different phenotypes of PD may be connected to polymorphism of fibrils, as it has been described earlier that the fibrils may have different subtypes (9). While the mature fibrils are known to be toxic for the cells, however, in recent years the intermediate species have been highlighted as even more neurotoxic (5, 15–17).

Many of the non-motor symptoms are associated with impaired peripheral nervous system or the peripheral part of the central nervous system (vagus nerve, olfactory bulb, etc.) (18, 19). The receptor neurons of the olfactory bulb are exposed directly to the environmental, giving an interface where environmental factors could trigger α-Syn aggregation (19). For α-Syn aggregation to occur, the enteric neurons (axons of myenteric plexus and/or submucosal plexus) needed to be triggered by an intrinsic or environmental factor (20). Recent study highlighted that enteroendocrine cells (EECs) were directly linked to enteric neurons, and therefore to the brain through the gastrointestinal muscles and the vagus nerve (21). Further, EECs also contained native α-Syn naturally, thus, these cells at the interface between environmental toxins and the enteric nervous system might be the source of the aggregation (21, 22). Several earlier studies supported a notion that the prion-like propagation of the α-Syn aggregation from cell to cell (19, 23–26). Interestingly, α-Syn aggregates also be found inside gastrointestinal nerves, from oesophagus to the rectal end, before they can be observed in any of the dopamine producing neurons (27). A direct transportation of α-Syn, injected into enteric neurons, towards the brain through the vagus nerve is demonstrated in several animal models, however, bidirectional propagation of α-Syn is possible (28–31).

Raman microspectroscopy is an ideal technology for the use in medical and biochemical studies because of a high sensitivity and marker-free application (12, 32). Raman spectra indicate changes of protein secondary structure based on specific peak shifts (11). Previously, Raman microspectroscopy was utilized in mouse model of AD for tau plaques from the murine brain (33, 34). Further, Raman spectroscopy was used for detection of α-Syn aggregations in vitro studies (9, 10). However, none of the findings described either human PD patient or rodent models to our knowledge. Thus, we hypothesized Raman microspectroscopy could be utilized to recognize α-Syn forms, either native or aggregations, and their current structure in the fibrillization process to gain an insight into the progression of PD from gut to brain.

In this study, we used BAC-SNCA transgenic rats (TG) expressing full length non-mutated human α-Syn (35) and control wild-type (WT) rats at different ages [4 months (4M) and 12M] to investigate the effects of ageing and the differences between normal and pathological tissues due to expression of human α-Syn in the colon. Using Raman imaging and microspectroscopy together with immunofluorescence staining we detected the presence of α-Syn aggregated proteins and confirmed changes in the protein secondary structures due to the fibrillization process in the brain and the colon tissues.

## Material and Methods

### Animals used for the study

The BAC-SNCA transgenic (TG) rats were described earlier (35) and corresponding age and sex matched wildtype (WT) rats (Sprague Dawley outbred genetic background) were used for this study. All the rats were kept in standard open type IV cages (3-4 rats/cage) under a 12 h light-dark cycle with *ad libitum* access to food pellet and water. All experiments were performed according to the EU Animals Scientific Procedures Act and the German law for the welfare of animals. All procedures were approved (TVA: HG3/18) by the authorities of the state of Baden-Württemberg.

### Colon and brain sample preparation

The brain and colon tissues of WT and TG rats aged 4M or 12M were used for this study. The tissues were frozen in O.C.T. compound and stored at −80°C until sectioning. Before sectioning tissues were acclimatized at −20 °C. A cryotome was used to cut tissue sections of 10 μm thickness, which were collected on standard glass slides. For the colon tissues 2-3 sections per slide were collected, while for the brain tissues one section per slide was collected. Eventually, 25 slides were collected for each sample. The remaining whole animal tissues were embedded into O.C.T. for protection and transported to the −80 °C freezer. The tissue sections were stored in the −20 °C freezer until further processing (details of chemical used in the study are available in Suppl. Table 1).

### Hematoxylin and Eosin staining

Hematoxylin and eosin staining (H&E) was performed on selected colon tissues (slide number 5, 19, 15, 20, 25) for morphological evaluation of the tissue sections and the identification of the area of interest. The tissue sections were washed three times with DPBS-for 5 minutes. Then, the sections were fixed with 4% PFA for 15 minutes and washed again with DPBS for 15 minutes. The sections were then treated with hematoxylin solution for 9 minutes and afterwards with demineralized water for a few seconds. A microscope was used to examine if the staining was sufficient. Next, the sections were left under running warm water for 10 minutes to wash away excessive staining and briefly washed with demineralized water for a few seconds. Afterwards, the sections were treated with eosin solution for 2 minutes, before being treated with demineralized water again for a few seconds. Then, the samples were put through a dehydration procedure of 70% ethanol (EtOH), 90% EtOH and 100% EtOH for 5 minutes each (details of chemical used in the study are available in Suppl. Table 1). Next, the sections were washed with isopropanol for 5 minutes twice. The sections were then mounted with isomount and covered with a thin glass slide. Afterwards the sections were scanned with the slide scanner.

### Nissl staining

Nissl staining was performed on selected brain tissues (slide number 5, 19, 15, 20, 25) for differentiation of the different brain regions, and to detect if the sections were intact enough for further processing. The sections were washed three times with DPBS for 5 minutes. If no more O.C.T. could be observed, the tissues were fixed with 4% PFA for 15 minutes and washed again with DPBS for 15 minutes. The sections were then treated with 1% cresyl violet solution for 10 minutes and afterwards with demineralized water for a few seconds (details of chemical used in the study are available in Suppl. Table 1). Next, a microscope was used to examine if the staining was sufficient. Afterwards, the samples were dehydrated analogously to H&E staining and washed twice with isopropanol for 5 minutes and mounted with isomount before being covered with a thin glass slide. Afterwards the sections were scanned with the slide scanner.

### Immunofluorescence staining

Immunofluorescence staining was performed at least once per samples so that the alpha-synuclein expressing regions could be identified. The primary and secondary antibodies were used for the staining (Suppl. Table 2). It was performed either with a single primary and a single secondary antibody or with two primary and two secondary antibodies.

The following procedure of antibody staining was modified slightly from the protocol previously established (36). The sections were placed into tubic racks and washed twice with DPBS for 10 minutes, and fixed with 4% PFA for 20 minutes, before being washed again with DPBS twice for 5 minutes. The unspecific binding sites were blocked with goat blocking buffer. Next, the samples were treated with primary antibodies for an hour. After a washing step with the washing buffer, the samples were treated with the secondary antibody for 30 minutes in a dark room at room temperature. Afterwards, another washing step was performed. If the samples were going to be measured directly with the Raman microspectrometer, the process was stopped, and the sections were stored in DPBS in a dark container for further use. If the samples were going to be just imaged with the fluorescence microscope, the samples were treated with DAPI for 10 minutes and after a washing step, the sections were mounted with prolong gold antifade mounting media (Thermofisher).

A control sample was always processed alongside the immunofluorescence staining for the evaluation of the staining success and the evaluation of unspecific background staining. The previous procedure was performed with the exception that instead of diluting the primary antibodies in the dilution buffer, the dilution buffer was used on its own. Afterwards the stained samples could be compared with the controls.

### Imaging of immunofluorescence-stained sections with observer microscope

The sections stained with DAPI were examined with the Observer fluorescence microscope (Zeiss GmbH, Germany). The 10x, 20x and 40x objectives were used, where with the 40x objective an immersion oil had to be applied to the samples. The microscopy was performed at a wavelength of 358 nm (DAPI, blue channel), 488 nm (green channel) and 594 nm (red channel). The software Zeiss Zen Blue Edition was used for the evaluation and processing of the images (Suppl. Table 3 and 4).

### Raman imaging and microspectroscopy

#### System set-up and sample preparation

A commercial Raman microspectroscope system (Alpha300R, WITec, Ulm) was used for all Raman measurements as previously described in detail (37, 38). The samples examined were either untreated colon samples or immunofluorescence-stained colon or brain samples. All samples were kept in DPBS before and during the Raman measurements.

First brain then colon samples were treated through immunofluorescence staining; therefore, three of the six measured areas were selected from stained regions while the other three were selected randomly from non-stained regions. The large area scan width to height was 50 μm x 50 μm, the points per line and lines per image of the scan were 100 and 100, making the scan step size 0.5 μm. The integration time was selected as 0.5 seconds. A fluorescence image and a bright field image of the same area were overlapped to identify the immunofluorescence-stained area. In addition, single spectra were measured for later analysis. 12 measurements were taken for the stained regions and 12 measurements for the non-stained regions within one sample, with 10 accumulations and an accumulation time of 10 seconds.

The untreated colon sections were measured by selecting three randomized areas in the muscularis externa. The large area scan width to height was 50 μm x 100 μm, the points per line and lines per image of the scan were 100 and 200, making the scan step size 0.5 μm. The integration time was selected as 0.5 seconds. The immunofluorescence-stained colon sections were measured with the same parameters, but stained regions were specifically selected for the measurements. A fluorescence image and the bright field image of the same sample area were overlapped to identify the stained area. Additionally, 15 single spectra were measured for each sample in stained regions for later analysis, with 10 accumulations and an accumulation time of 10 seconds.

#### Pre-Treatment of the Spectra

For later statistical analysis, the spectra were pre-treated with the Project Five software (WITec, Ulm, Germany; Suppl. Table 4). Cosmic rays were removed. Next, the background was subtracted from the spectra using a shape correction method and a shape of 150. Therefore, the baseline was similar in all spectra and therefore comparable. The spectra were then normalized, with the normalization type area to 1. Large area scans were stitched together so later they could be analyzed together.

#### Raman imaging and spectral analysis

Raman data were analyzed through different methods to detect differences in the samples at different time-points (4 months or 12 months) or at a different modification (TG or WT).

The spectra (overall average of the Raman images of all the samples in each group) of the different groups (4M WT & TG and 12M WT & TG) were then compared to each other to identify differences. The ratio of different peaks from amide I and amide III bands were compared. The compared peaks were phenylalanine (1004 cm-1)/amide III – β-sheet (1267 cm-1), phenylalanine (1004 cm-1)/amide III – α-helix (1298 cm-1), phenylalanine (1004 cm-1)/amide III – α-helix (1340 cm-1), phenylalanine (1004 cm-1)/amide I – α-helix (1658 cm-1), amide III – β-sheet (1267 cm-1)/amide III – α-helix (1298cm-1), amide III – β-sheet (1267 cm-1)/amide III – α-helix (1340cm-1) and amide III – β-sheet (1267 cm-1)/amide I – α-helix (1658cm-1). The statistical analysis through t-test was performed with Microsoft Excel 365.

#### Principal component analysis (PCA)

Single spectra data were uploaded to Unscrambler X 10.5 to perform principal component analysis (PCA). The different sample groups were analyzed with seven principal components (PC) as described previously (37, 38). The results were presented in a score plot explained by two PCs. For each PC, a loading plot was obtained, where the scores were explained through spectra. The results from different groups were compared statistically through t-test with Microsoft Excel 365. The PCs with statistical significance and best separation were selected.

#### True component analysis (TCA)

True component analysis (TCA, Project Plus Software, WITec) was used to analyse Raman images as previously described in detail (37, 38). For the colon samples, the analysis was performed in the whole area (intensity range of the pixels: 0-1) or in the stained regions (intensity range of the pixels: 0-0.8).

#### Statistics

For spectral analysis, PCA and TCA Raman images/scans were used (Suppl. Fig. 1a, b and Suppl. Table 4). The samples were measured and separated into four group: 4M WT, 4M TG, 12M WT and 12M TG. The single spectra of the different groups were compared to each other to identify the difference between sample groups. Further, the intensity and the full width at half maximum (FWHM) of the spectra were statistically analysed based on a t-test or one-way analysis of variance (ANOVA).

## Results

### Identification of endogenous α-Syn aggregation in the brain olfactory bulb region of TG rats

To identify the accumulation of α-Syn aggregation in the TG brain, we used the olfactory bulb brain regions as it is an early site of α-Syn accumulation (39, 40). The brain sections were used for Nissl staining for the confirmation that the area of interest was suitable for the Raman measurements (Suppl. Fig. 1a, b). We detected endogenous rat-specific α-Syn faintly in the whole brain region of either WT or TG but predominantly on the edges (Fig. 1a). The α-Syn stained area was used for the measurement by Raman microspectroscopy (Suppl. Fig. 1b). After measurement of the samples, the assignment of the peaks was identified for comparable results found in the literature (Suppl. Fig. 1c, Suppl. Table 5).

**Figure 1.**
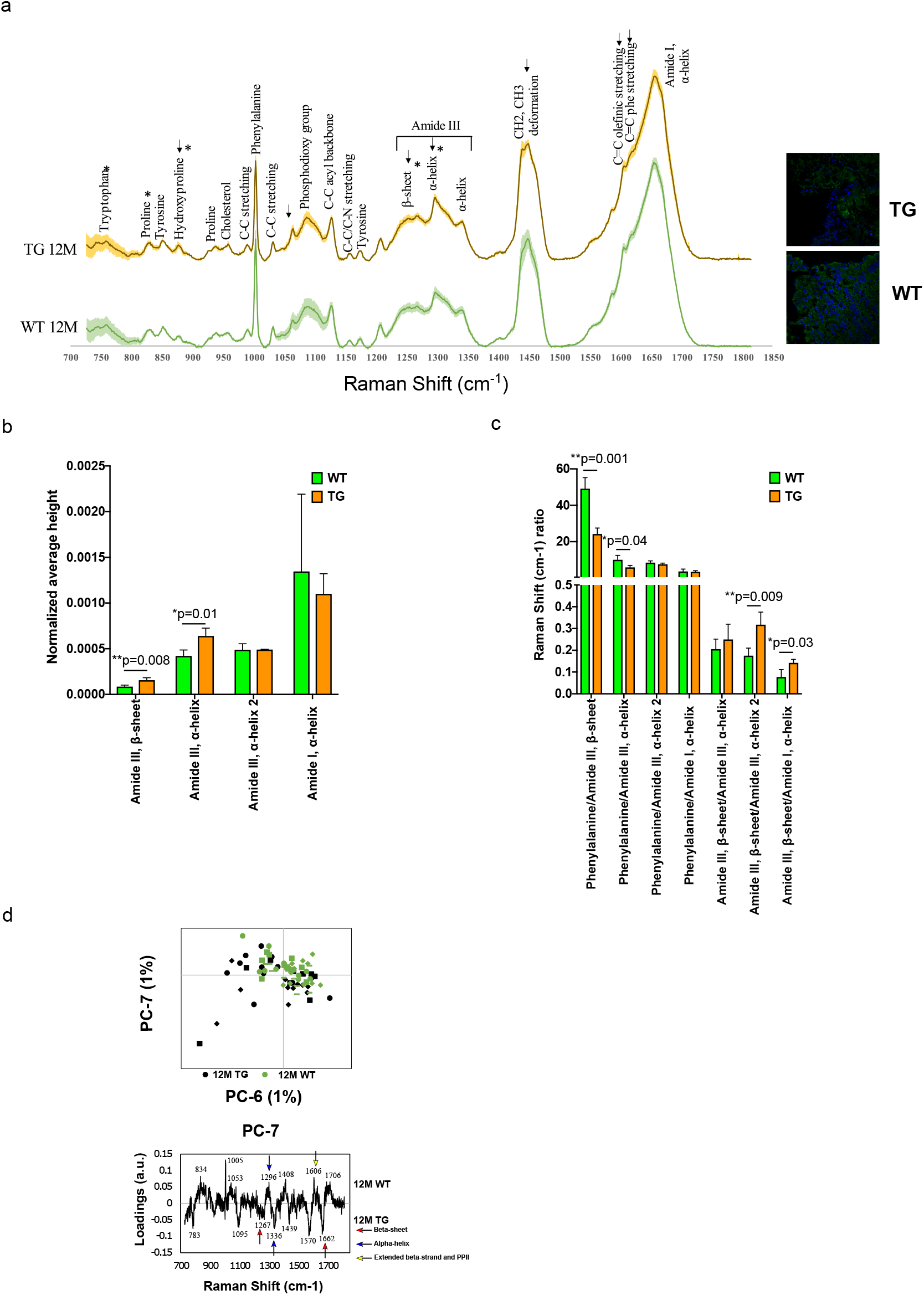
Spectral comparison in 12M WT and TG olfactory bulb regions. (a) The spectra of WT and TG olfactory bulb regions. (b) All the marked changes were from the statistical 12M WT and TG intensity comparison based on Student’s unpaired t-test. Differences were detected in the 759 cm-1 (p=0.04), 830 cm-1 (p=0.02), 877 cm-1 (p=0.04), 1268 cm-1 (p=0.008) and 1298 cm-1 (p=0.01) peaks. P value represents *(p ≤0.05), **(p ≤0.01). (c) The ratio of different peaks from amide I and amide III were compared with Student’s unpaired t-test to identify statistical changes. Differences were detected in the phenylalanine/amide III – β-sheet (p=0.001), phenylalanine/amide III – α-helix (p=0.04), amide III – β-sheet/amide III – α-helix (p=0.009) and amide III – β-sheet/amide I – α-helix (p=0.03). P value represents *(p ≤0.05), **(p ≤0.01). (d) Comparison of genotype (12M WT and TG) samples through PCA with scores and loadings for the brain sample. The 12M WT and TG 12M samples were compared with PCA of PC-6 and PC-7. Both PC-6 (p=0.03) and PC-7 (p=0.006) were significant (upper panel). The loadings of PC-6 were visualized (lower panel). Positive side WT while negative side TG brain samples.

First, we compared Raman spectra of WT and TG genotypes at 12M age. We can observe the visible differences for the peaks 879 cm^-1^ (hydroxyproline), 1063 cm^-1^ (C-C skeletal stretch), 1265 cm^-1^ (amide III, β-sheet), 1298 cm^-1^ (amide III, α-helix), 1450 cm^-1^ (CH_2_, CH_3_ deformation) and the amide I shoulders 1586 cm^-1^ (C=C olefinic stretch) and 1606 cm^-1^ (C=C phenylalanine stretch). Based on all the biological samples (n=3) for 12M TG rat brains had a higher intensity than the 12M WT rat brains (n=4) except on the amide I shoulders, where WT rat brains have more intensity (Fig. 1a). At 1450 cm^-1^, the peak produced a shoulder at 1441 cm^-1^ in the 12M TG rat brains, while it was totally absent in the 12M WT rat brains, this could be due to polarization effect. We found that intensity of the 759 cm^-1^, 830 cm^-1^, 877 cm^-1^, 1268 cm^-1^ (amide III, β-sheet) and 1298 cm^-1^ (amide III, α-helix) peaks were significantly different between WT and TG rat brains (Fig. 1b, Suppl. Fig. 2). Significant differences were present when spectral peak data were normalized in the 1004 cm^-1^ (phenylalanine) to 1267 cm^-1^ (amide III, β-sheet) ratio and the 1004 cm^-1^ (phenylalanine) to 1298 cm^-1^ (amide III, α-helix) ratio (Fig. 1c). Further, significant difference was also noticed when spectral peak data were normalized in the 1267 cm^-1^ (amide III, β-sheet) to 1340 cm^-1^ (amide III, α-helix) ratio and the 1267 cm^-1^ (amide III, β-sheet) to 1658 cm^-1^ (amide I, α-helix) ratio (Fig. 1c).

### Genotype comparison for PCA

The Raman spectra of 12M WT & TG brains were compared to each other through PCA to identify differences between the genotypes. A statistically significant separation was achieved in the fingerprint region of the PC-6 and PC-7 score values (both PCs explained 1% of the variance) (Fig. 1d). It was possible to detect the separation as most of the WT 12M samples were on the positive region, while most of the TG 12M samples were on the negative region (Fig. 1d, upper panel). The prominent peaks present on PC-7 and PC-6 loadings were shown in Fig. 1d, lower panel and Suppl. Fig. 3 respectively. Two Beta sheets were present in 12M TG rats compared with WT, while 12M WT rats have more alpha helix and beta-stranded and PP II than 12M TG rat brain.

### Brain genotype comparison using Raman imaging

The area of interest for Raman imaging was identified *via* α-Syn immunofluorescence staining. In Raman images from all samples four major spectral components were identified: lipids, cell nuclei, matrix and an unknown component (Fig. 2). The measured representative regions from 12M WT and TG along with the separated components are shown in Fig. 2b. No significant difference was observed in cell nuclei and unknown component except some visible difference in matrix and significant difference in lipids (Fig. 3). Lipids appeared to be more solid in the TG rat brain with less space in between than the WT brain samples (Figure 2c). Further, in the lipid component, intensity statistical significant differences were detected at 873 and 1175 cm^-1^ (Tyrosine) between 12M WT compared with 12M TG (Fig. 3 a,c). However, some visible differences were also observed 1439 and1083 cm^-1^, though it did not reach to a significance level (Fig. 3 a,c). Similarly, matrix component in the brain region was appeared to different at 1209, 1589 (C=C olefinic stretching) cm-1 peaks and some visibly different (Fig. 3 b,d). Taken together, we concluded that TG rat brain samples have different property compared to WT brain samples.

**Figure 2.**
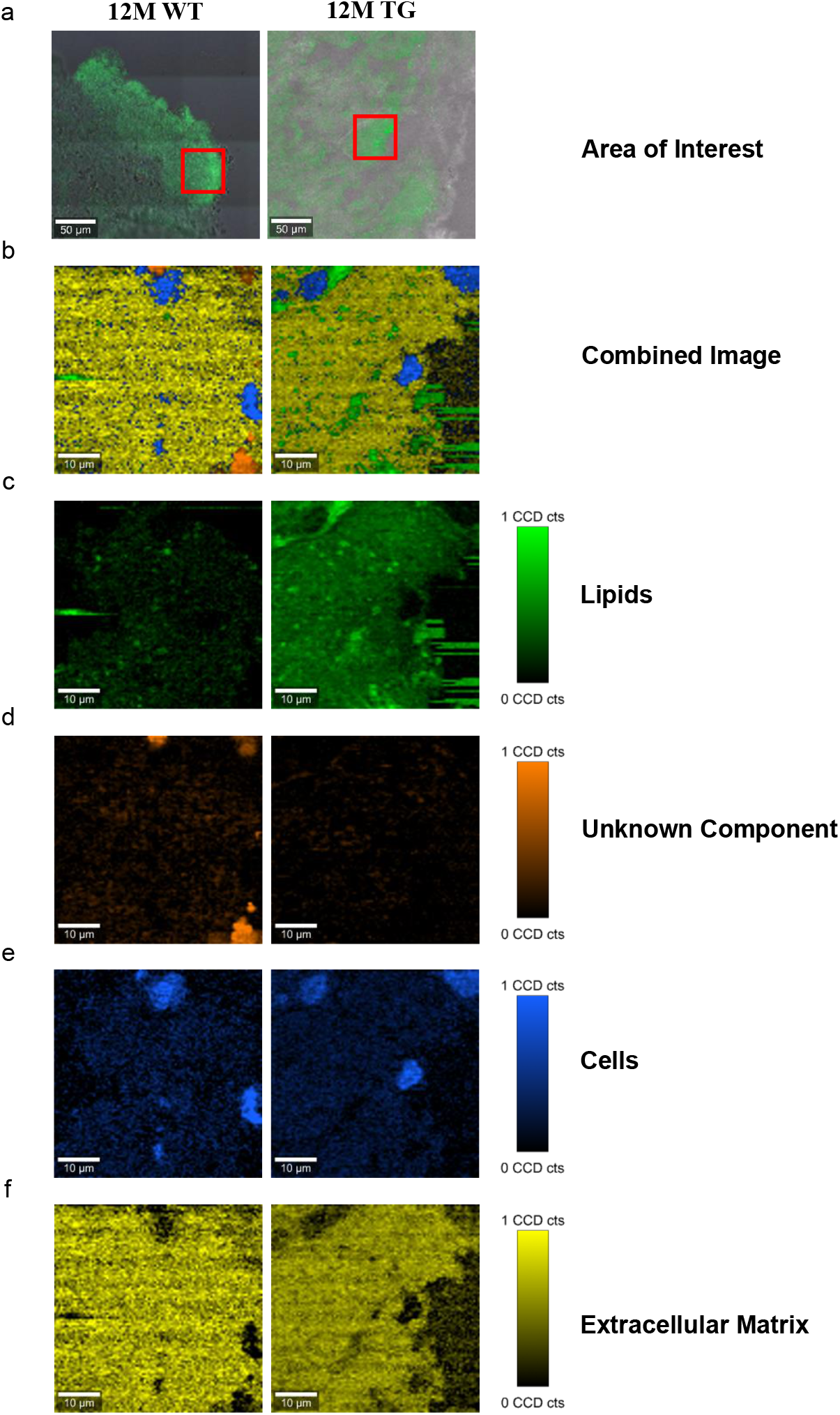
TCA of selected areas in the olfactory bulb of WT and TG animals. (a) The measured area was shown in red boxes. (b) The combined image of the components was shown. (c) The separated components through TCA were lipids in green, (d) an unknown component in orange, (e) cells in blue, and (f) extracellular matrix components in yellow. The CCD count interval was listed for all the components.

**Figure 3.**
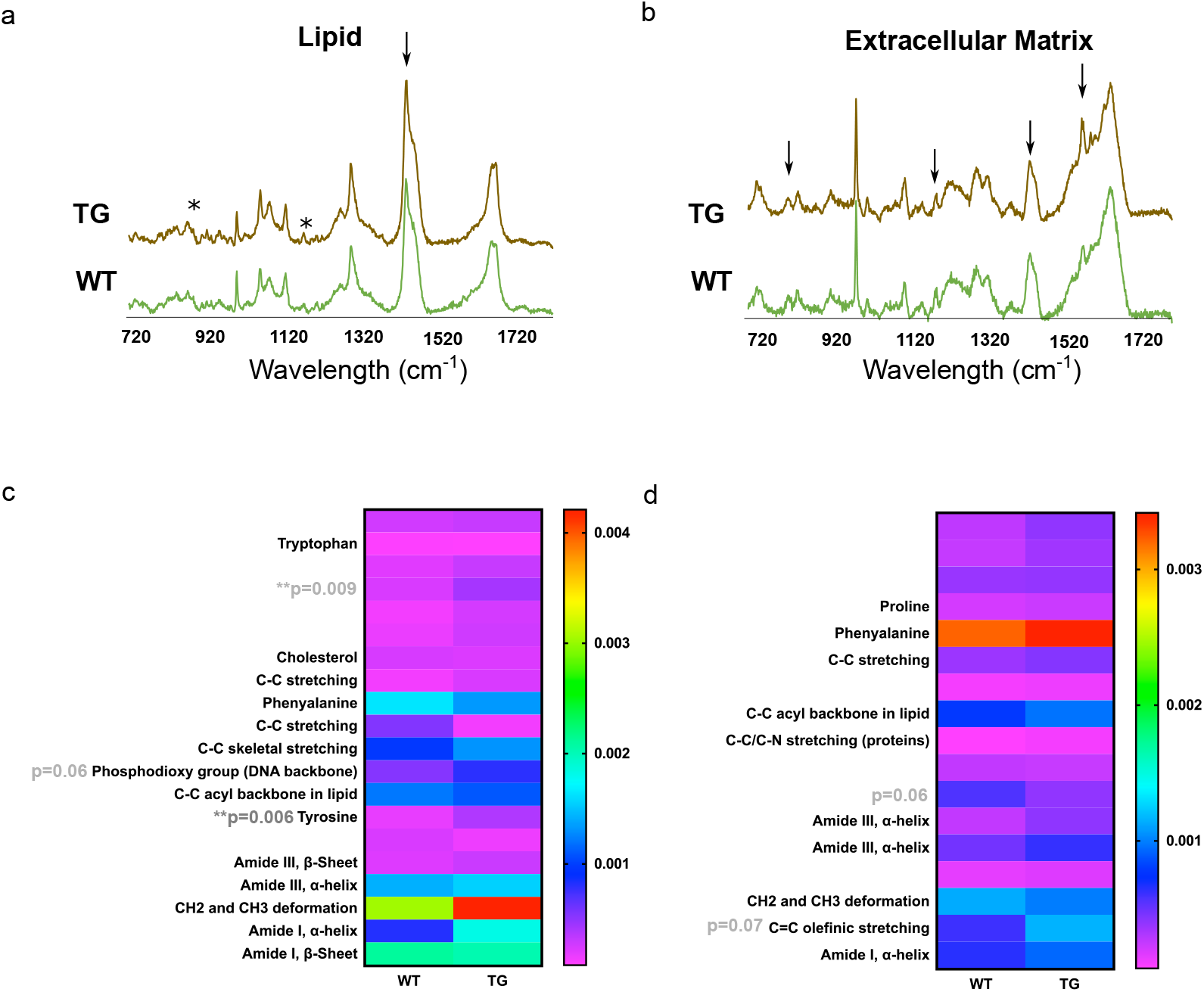
Estimation of lipids and extracellular matrix components in the olfactory bulb region. The components displaying the same results were put together and averaged in their groups. The two components (lipids and extracellular matrix) out of five were appeared to be different in most samples. (a) lipids and (b) extracellular matrix component spectra from WT and TG brain olfactory bulb region. X-axis represents the wavelength in cm^-1^ while y-axis shows the spectral intensity. The separated groups were 12M WT (green) and 12M TG (brown). (c) Differences in the lipid component were observed for intensity at 873 cm^-1^ (p=0.009) and 1175 (p=0.006) between 12M WT and TG based on Student’s unpaired t-test and data shown as a Heatmap. (d) Heatmap represents the extracellular matrix component, some apparent difference in the spectra; however, it did not reach to a significance level. P value represents *(p ≤0.05), **(p ≤0.01).

#### Identification of endogenous α-Syn aggregation in the colon of TG rats

After establishing the spectral pattern in the brain, we focussed on the colon region with the predicted α-Syn aggregations could be present in TG rats. After the measurement of the samples with the Raman micro-spectroscopy, the assignments of the peaks were identified for comparable results as described in the brain samples (Fig. 4a). For spectral analysis and PCA, single spectra were measured while for TCA, large area scans were used (Suppl. Fig. 1c, d).

**Figure 4.**
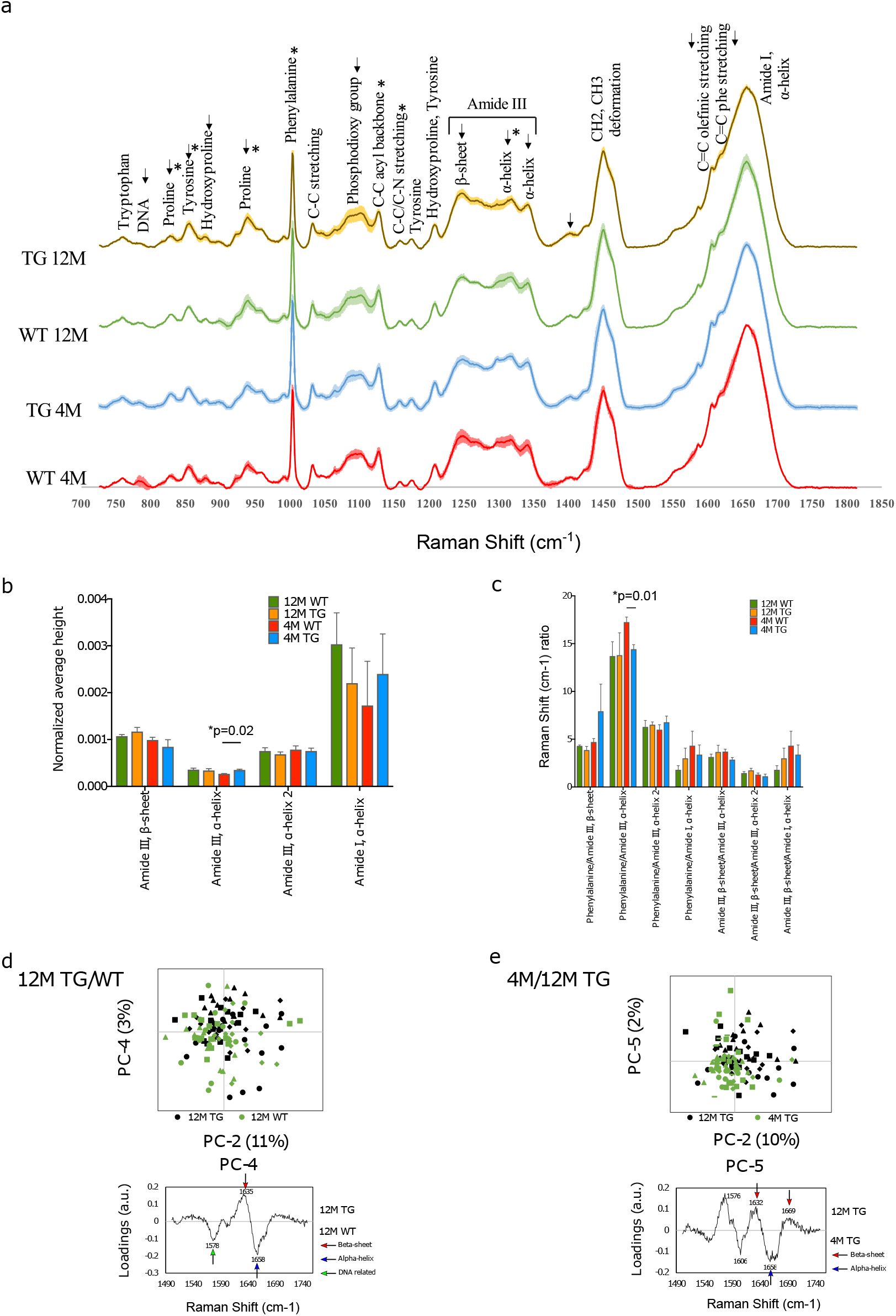
Spectral comparison at the genotype and age of the WT and TG rat colons. (a) The spectra of WT and TG colon samples at 4 and 12M age. (b) Differences were detected at the intensity at 4M age for WT and TG in the 1317 cm-1 (p=0.01) peak. (c) The ratio of different peaks from amide I and amide III were compared with Student’ unpaired t-test to identify statistical changes. Differences were detected in the phenylalanine/amide III – α-helix ratio (p=0.01). P value represents *(p ≤0.05). (d) Comparison of the amide I 12M TG vs. WT samples through PCA with scores and loadings in colon. Amide I samples were compared with PCA at PC-2 and PC-4 (upper panel). PC-4 was significant (p=0.01). The loadings of amide I PC-4 were visualized (lower panel). (e) Comparison of the amide I between 4M and 12M TG samples through PCA with scores and loadings in colon. Both PC-2 (p=0.00003) and PC-5 (p=0.004) were significant (upper panel). The loadings of amide I, PC-5 were visualized (lower panel).

First, we made the genotype comparisons between 4M WT and TG colon samples using spectral analysis (Fig. 4b). A clear visible difference was identified in 879 cm^-1^ (hydroxyproline), 1093 cm^-1^ (phosphodioxy group), 1248 cm^-1^ (amide III, β-Sheet), 1317 cm^-1^ (amide III, α-helix) and 1342 cm^-1^ (amide III, α-helix). Only, the intensity of the 1317 cm^-1^ (amide III, α-helix) peak was statistical different between 4M WT and TG colon (Fig. 4b). The chosen peaks were from amide I or amide III divided with each other and also to phenylalanine (1004 cm^-1^), significant differences were detected at 1003 cm^-1^ (phenylalanine) to 1317 cm^-1^ (amide III, α-helix) ratio (Fig. 4c). Further, 12M WT and TG samples were analysed. Visible differences were observed in the 855 cm^-1^ (tyrosine), 879 cm^-1^ (hydroxyproline), 938 cm^-1^ (proline), 1096 cm^-1^ (phosphodioxy group), 1248 cm^-1^ (amide III, β-Sheet), 1317 cm^-1^ (amide III, α-helix) and 1402 cm^-1^ (C=O stretch) (Fig. 4a). The 12M TG colon samples were more intense in every marked area except the 1317 cm^-1^ peak. However, statistically significant differences were observed in the FWHM of the 1128 cm^-1^ peak (Suppl. Fig. 4a).

Further ageing comparisons were made among 4M and 12M TG colon samples. The 12M TG colon sample peaks were more intense in every marked area except the 827 cm^-1^ peak and the two α-helix peaks of amide III (1299 cm^-1^ and 1342 cm^-1^) where, as identified in previous sections, a change in intensity was observed at the β-sheet to α-helix turning point. Statistically differences were observed in the intensities of the 829 cm^-1^, 855 cm^-1^, 939 cm^-1^, 1004 cm^-1^ and 1158 cm^-1^ peaks (Suppl. Fig. 4b). No statistical difference was observed in 4M and 12M WT rat colon samples (Fig. 4a,c). Thus, our spectral data highlight that ageing could have an important change in the colon tissue composition of the TG rats based on peak intensities.

### PCA analysis to identify the aggregation of *α-Syn* in the TG colons

The 4M TG/WT samples were compared to each other through PCA to identify the differences through the modification of rats at the same age. The PCA of the fingerprint region was analysed. Significant separation was observed at the PC-4 score values, which explained 8% of the variance, as the other PCs did not show differences. The separation at PC-4 was easily distinguishable, with the WT 4M samples mostly on the negative region and the TG 4M samples mostly on the positive region (Suppl. Fig. 4c). The most prominent peaks were 1001 cm-1 (phenylalanine) and 1321 cm-1 (amide III α-helix) on the positive region (4M TG) and 787 cm-1 (DNA) and 1378 cm-1 (T, A, G) on the negative region (4M WT). Most of the major peaks in the negative region were related to DNA.

Furthermore, in amide I, a significant separation was achieved through the PC-2 score values, which explained 8% of the variance and the PC-7 score values, which explained <1% of the variance (Suppl. Fig. 4d). The loadings for PC-2 showed the peaks 1637 cm-1 (β sheet) and 1685 cm-1 (turn) on the positive region and 1586 cm-1 (C=C olefinic stretch) and 1606 cm-1 (C=C phenylalanine stretch) on the negative region. The loadings for PC-7 showed the peaks 1630 cm-1 (β sheet) and 1654 cm-1 (α-helix) on the positive region and 1638 cm-1 (β sheet) and 1670 cm-1 (extended β-strand and polyproline II (PPII) structures) on the negative region. In amide III, a significant separation was achieved through PC-1 score values, which explained 41% of the variance and PC-4 score values, which explained 7% of the variance (Suppl. Fig. 4d). The loadings at PC-1 showed the peaks 1294 cm-1 (α-helix) and 1340 cm-1 (α-helix) on the positive region and the peak 1245 cm-1 (β sheet) with the shoulder 1270 cm-1 (α-helix) on the negative region.

The 12M TG and WT samples were compare to each other through PCA to identify differences of modified rats at the same age (12M). PCA of fingerprint region did not show any difference among WT and TG rat samples. The amide I and amide III regions were analysed closer with separate PCAs. In amide III, a significant separation was achieved through the PC-6 score values, which explained 1% of the variance. The loadings of PC-6 showed 1262 cm-1 (β-sheet) and 1310 cm-1 (α-helix) on the positive region and 1296 cm-1 (α-helix) on the negative region (Suppl. Fig. 5a). In amide I, a significant separation was achieved through the PC-4 score values, which explained 3% of the variance (Fig. 4d). The separation at PC-4 was overlapping, but recognizable. The 12M TG samples were more on the positive region, while the 12M WT samples were more on the negative region (Fig. 4d). The most significant PC-4 was visualized with PC-2 with their corresponding loadings (Fig. 4d). The loadings of PC-4 showed the peak 1635 cm-1 (β-sheet) on the positive region and the peaks 1578 cm-1 (nucleic acids) and 1658 cm-1 (α-helix).

Further, ageing comparisons were made in TG rats (4M vs 12M) samples through PCA to identify the differences among TG colon samples with ageing. Significant separation was detected at the PC-2 score values (25% of the variance), at the PC-3 score values (15% of the variance) and at the PC-4 score values (11% of the variance) (Suppl. Fig. 5b). The most significant PC-4 was visualized with PC-2 with their corresponding loadings (Suppl. Fig. 5b). 12M TG samples were more on the positive region and the TG 4M samples were more on the negative region. The separation at PC-2 was less clear but distinguishable with the TG 12M samples more on the positive region and the TG 4M samples more on the negative region. The major peaks on PC-2 were 860 cm^-1^ (phosphate group), 940 cm^-1^ (proline), 1248 cm^-1^ (amide III, β-sheet), 1653 cm^-1^ (amide III, α-helix) and 1685 cm^-1^ (amide I, turn) on the positive region and 1065 cm^-1^ (C-C skeletal stretch), 1086 cm^-1^ (phosphodioxy group), 1130 cm^-1^ (C-C acyl backbone), 1296 cm^-1^ (amide III, α-helix) and 1439 cm^-1^ (CH_2_ and CH_3_ deformation) on the negative region. The loadings of PC-4 contained the main peaks 1637 cm^-1^ (β sheet) and 1673 cm^-1^ (extended β-strand and PPII structures) on the positive region and the peaks 1001 cm^-1^ (phenylalanine) and 1315 cm^-1^ (amide III, α-helix) and on the negative region.

The amide I and amide III regions were analysed further in detail with separate PCAs (Suppl. Fig. 5c). In amide I, a significant separation was achieved through the PC-2 score values, which explained 10% of the variance, the PC-3 score values, which explained 6% of the variance and the PC-5 score values, which explained 2% of the variance (Fig. 4e). The separation was clearer with PC-5, where the TG 12M samples were more on the positive region, while the TG 4M samples were mostly in the negative region. The loadings for PC-2 showed the major peaks 1634 cm-1 (β sheet), 1668 cm-1 (β-sheet) and 1683 cm-1 (turn) on the positive region and the peak 1605 cm-1 (C=C phenylalanine stretch) on the negative region (Suppl. Fig. 5c). The loadings for PC-5 showed the major peaks 1576 cm-1 (nucleic acids), 1632 cm-1 (β sheet) and 1669 cm-1 (β-sheet) on the positive region and 1606 cm-1 (C=C phenylalanine stretch) and 1658 cm-1 (α-helix) on the negative region (Fig. 4e). In summary, 12M TG contained higher amount of β-sheet compared with 4M TG, while 4M TG contained advanced intermediate oligomers and more α-helices.

### TCA analysis to detect other cell components with ageing in TG rat colon

Every measured sample with Raman microspectroscopy was also subjected to TCA. We stained the colon tissues with α-Syn staining (D37A6) antibody to achieve the proper results for the TCA and only antibody positive stained region was used. (Fig. 5a, b).

**Figure 5.**
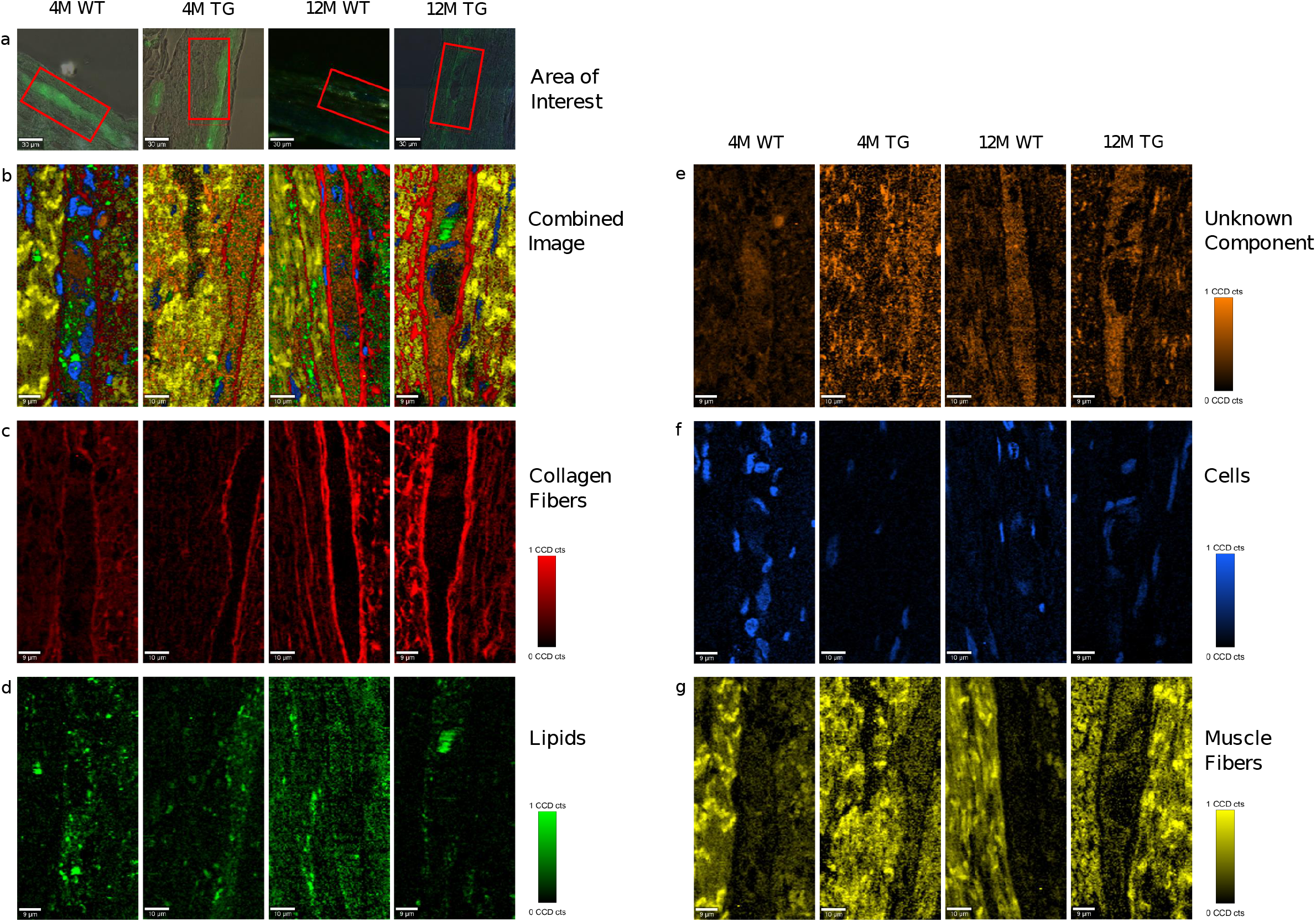
TCA of selected areas in the colon of WT and TG rats. (A) The measured area was shown in red boxes. (B) The combined image of the components was shown. (C) The separated components through TCA were collagen fibers in red, (D) lipids in green, (E) unknown component in orange, (F) cells in blue and (G) muscle fibers in yellow. The CCD count interval was listed for all the components.

Raman images showed that collagen fibers surrounded the α-Syn-stained regions in all the samples with a more intense appearance in the older rats, presumably increasing in thickness with age (Fig. 5c). Lipids appeared to be more concentrated in the stained regions with no changes in intensities between genotypes (Fig. 5d). The colon unknown component appeared more solid in the α-Syn stained regions, possibly showing a connection with the protein. The intensity was lower in the WT 4M region, with no visible changes in the other groups (Fig. 5e). The cells were mostly observed in the non-stained regions in all animals except the 4M WT sample, where cells were also observed inside the stained area (Fig. 5f). The cells in the TG groups appeared smaller, possibly in the process of apoptosis. The muscle fibers were concentrated in the regions surrounding the staining, with no visible changes in intensity between the groups (Fig. 5g). The spectral components of TCA were separated into their respective groups (4M WT, 4M TG, 12M WT and 12M TG). The spectra particularizing the same component were averaged and graphed together so that the differences of the groups in the same component could be visualized (Suppl. Fig. 6).

After the extraction of the different components through TCA (collagen fibers, lipids, unknown component, cells and muscle fibers) the CCD counts and the number of pixels were scaled to an interval where all the positive pixels were contained. From the CCD count and the pixels, an intensity was calculated. The averaged intensity per pixel was taken for each component in each group to detect any difference between the components when their intensity in the TCA was compared. After statistical analysis, changes were observed in the muscle fibers between 4M and 12M TG as well as nearly significant among 4M WT and TG colon samples (Suppl. Fig. 6). Other components were not significantly different (data not shown).

Furthermore, the averaged intensities per pixel were taken for the components stained with α-Syn antibody (D37A6), in the regions where the staining was observed. A significant difference was also observed at 4M age among WT and TG for collagen fibers (Fig. 6a-b).

**Figure 6.**
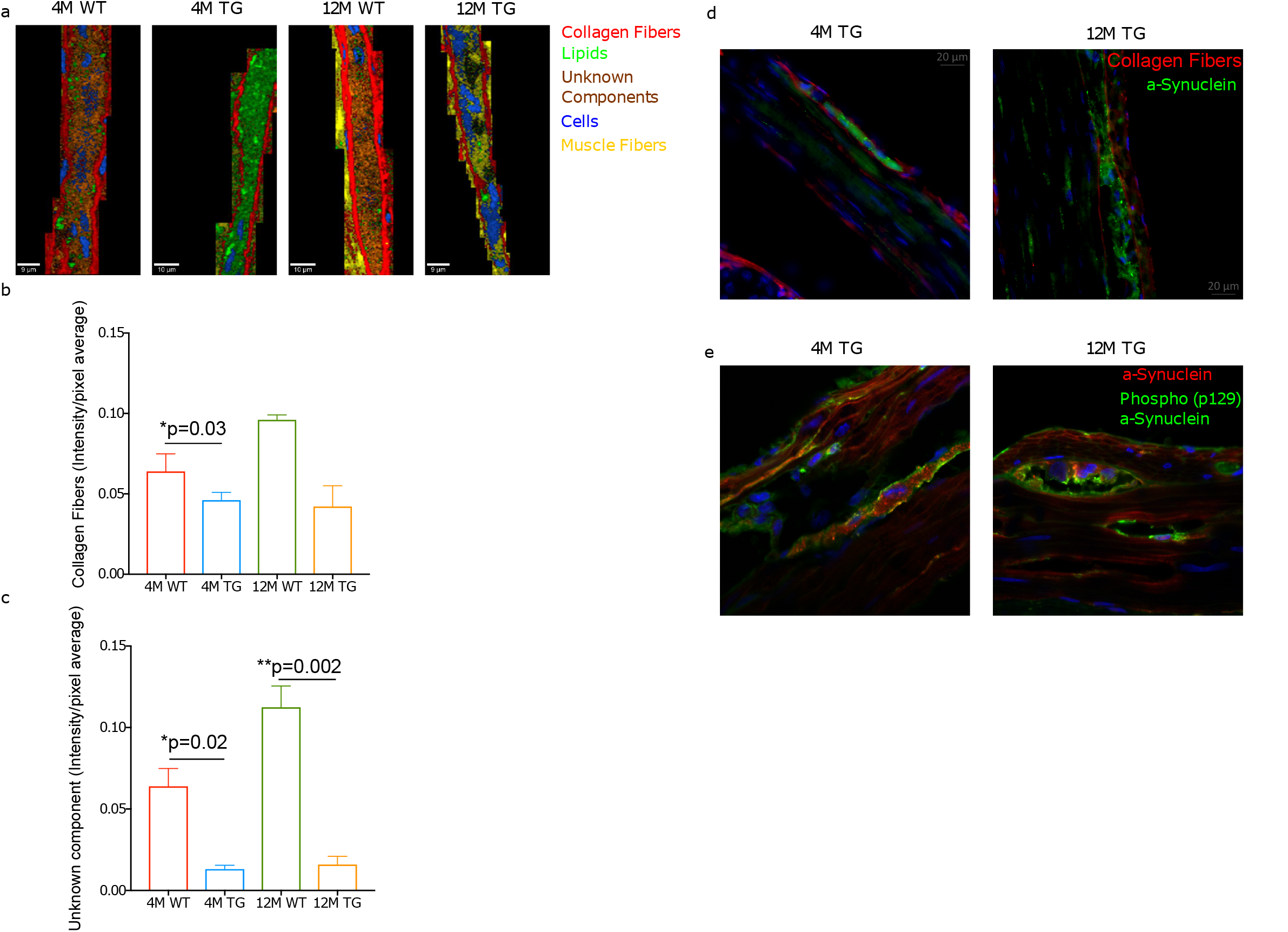
Collagen fibers and an unknown component in the TG rat colon. (a) Combined representative TCA images of the stained regions. The components were collagen fibers (red), lipids (green), an unknown component (orange), cells (blue) and muscle fibers (yellow). The intensity range was scaled 0-0.8 for all components. (b,c) The bar diagram showed the collagen intensity/pixel average for WT and TG rat colon samples at 4M and 12M age. Averaged intensities per pixel of the collagen fibers and unknown component statistically compared using Student’s unpaired t-test. P value represents *p ≤0.05. Significant differences were observed in the unknown component between 12M WT and TG (p=0.002) as well as between 4M WT and TG 4M (p=0.03). In contrary, statistical difference was observed in the collagen fibers only for 4M WT and TG (p=0.02). P value represents *(p ≤0.05), **(p ≤0.01). (d) Staining of the colon tissues with α-Synuclein (green, D37A6), collagen I (red, 113M4774) antibodies and DAPI for cell nucleus for 4M and 12M TG rats. (e) The detection of phospho-α-Synuclein (p129; green) and α-Synuclein (red, D37A6) and DAPI (blue) in the colon tissues of 4M and 12M TG rats.

Additionally, statistical differences were observed at the unknown component between 12M WT and TG as well as for 4M WT and TG colon samples (Fig. 6c).

Further, the fluorescence microscope was used to display stained tissue sections for α-Syn and collagen fibers surrounding the α-Syn-stained regions (Fig. 6d). In 12M TG, pockets between the α-Syn and collagen fibers were observed (Fig. 6d). Cells (based on DAPI staining) were abundant in the surrounding regions but absent in the stained regions, occurring between adjacent collagen fibers shielding (Fig. 6d). In the 4M TG rats no unstained areas were observed between the α-Syn and the collagen fibers. Morphologically, cells differed from the surrounding cells outside the stained region, being smaller in size and shorter in length (Fig. 6d). Further, we stained the TG colon tissues using phospho-α-Syn antibody along with total alpha-Syn protein. This analysis revealed that in 12M TG colon tissue phosphorylation/aggregation is mostly located in the pocket region compared with 4M TG rat colon samples (Fig. 6e). Thus, explains that pocket region between the total α-Syn and collagen regions could be due to aggregated α-Syn in older TG rats. Overall, our Raman imaging and microspectroscopy data revealed that ageing could be involved in aggregation of α-Syn in the colon TG rats.

## Discussion

Based on the notion that the beginning of the aggregation process in the colon, α-Syn protein molecules infect the neurons in a prion-like manner, propagating up the vagus nerve towards midbrain, where it causes PD (5, 28, 30, 41). Therefore, aggregation of the proteins and changes in protein conformation was expected in younger rats in the colon, while a progressed disease was probable in older TG rats. We confirmed the presence of aggregated proteins, detected changes in the secondary structure of the proteins due to the fibrillization process, and identified the changes in colon tissues through a combination of Raman imaging and Raman microspectroscopy.

Our brain data suggested that signs of the fibrillization process was detected TG rats. Furthermore, the 12M TG rats contained dominantly β-sheet rich secondary structures, signs of an advanced PD with protofibrils and mature fibrils. TCA analysis further revealed changes in lipid structures. This was significant as native α-Syn is known to bind naturally to lipids, making lipid molecules a reflection of α-Syn molecules. Changes in the secondary structures of lipids indicate an effect of α-Syn aggregation and points towards native α-Syn being present in the TG rats.

While we used the olfactory bulb region in the brain (42) as the control against colon for our experiments, we believe it is still an acceptable control for the progression of PD in the brain as a 90% correlation between Lewy pathology in the olfactory bulb and *substantia nigra* in midbrain was described (43). Previously, it was detected behavioural difference reminiscent of progressive PD, with an early alteration of olfaction already in 3M TG rats (35). Olfactory bulb was easy to differentiate with other brain regions clearly. Further, the direct connection of the olfactory bulb with the midbrain regions also contribute to similar progression of pathology between the brain regions, making the olfactory bulb a reflection of the *substantia nigra* in most instances.

In colon, 4M TG rats showed a more advanced aggregation than brain as β-sheets were detected in higher numbers. The presence of additional α-helices and extended β-strands and PPII structures pointed towards an early fibrillization process, making protofibrils unlikely. Additionally, 12M TG rats on the other hand were rich in β-sheets, indicating advanced PD in the colon, could reflect the non-motor symptoms. TCA analysis and fluorescence microscopy images revealed that collagen fibers were surrounding the α-Syn stained regions. Averaged intensity/pixel showed that WT rats contained more collagen fibers, whereas in 12M TG rats the collagen fibers were likely destroyed due to inflammatory mediatory or direct effects of α-Syn. The unknown component of colon was present in α-Syn containing regions of WT rats, linking it to the native protein as well as cell death which was detected in the 12M TG rats.

No compelling signs of aggregation were observed in the colon in rats of any age in the WT samples, while less advanced fibrillization, possibly misfolded monomers or early spheroidal oligomers were detected in 12M WT samples in the brain, α-Syn aggregation could begin in advanced ages even in healthy individuals with a starting point of olfactory bulb, but not colon. As previously described (44), the failure of the UPS and LAS pathways are a likely cause of α-Syn aggregation in 12M WT rats. The lack of fibrillization in the colon support this theory, as the failure of UPS and LAS systems were only described in the brain but not in the enteric nervous system, indicating environmental factors as the more likely trigger in a-Syn aggregation in the colon such as gut bacterial dysbiosis (45). In the 4M TG group, the colon showed signs of a more advance fibrillization than brain, where a higher number of α-helices were detected, supporting several researchers, who illustrated the colon to brain path of the aggregation through the vagus nerve (27, 28, 30). These results confirmed the prion-like propagation hypothesized by Braak *et. al.(46)* with the enteroendocrine cells as the possible route between the lumen and the enteric nervous system (21).

The results were able to successfully confirm the presence of α-Syn aggregations in the colon enteric nervous system. Additionally, a more advanced fibrillization process was identified in colon of 4M TG rats, confirming the hypothesis of this study, where the fibrillization process starts in the colon before advancing towards the brain. In 12M older rats both regions showed signs of advanced fibrillization. Nevertheless, the limitations of the study should be noted for the next experiments, making the results possibly clearer. This is especially true for the antibody staining, where human aggregation specific α-Syn antibody would have been preferred. It should be noted that according to literature, the intermediate oligomers between monomers and fibrils were suspected to be the toxic agent, rather than the mature fibrils, marking the importance of study into the intermediate species, and priority of reversing the fibrillization process rather than slowing it down. The 4M TG group, where most of the intermediate species were present might therefore be in the period of most neuronal damage.

Future studies are required to understand all the unknown aspects. More time points (early or late stages of disease) in the rats would be of advantage to pinpoint the starting point of the aggregations more accurately. The measurement of several more brain regions, or even the vagus nerve, would be necessary to understand the further progression of the disease. As previously mentioned, aggregation-specific α-Syn antibodies and a larger sample size would have given more certain results.

In the future, Raman microspectroscopy could be a routine tool to detect PD disease in advance through analysis of colon biopsies with a considerable reduced misdiagnosis rate, leading to better care and life quality of the diseased.

## Declarations

### Ethics statement and approval

All the experiments were performed according to the EU Animals Scientific Procedures Act (2010/63/EU) and the German law for the welfare of the animals. All the procedure and methods were approved by the local government authorities (Regierungspräsidium, Tübingen; TVA HG3/18) of the state of Baden-Württemberg, Germany.

### Consent for publication

No patients or human data used in this study. All authors read the manuscript and approved to be co-authors on the manuscript and have substantial contribution in the manuscript.

### Availability of data and material

The datasets used and/or analysed during the current study are available from the corresponding authors on a reasonable request.

## Acknowledgements

We acknowledge support by Deutsche Forschungsgemeinschaft (DFG) and Open Access Publishing Fund of the University of Tübingen.

## Author’s contribution

EMB, YS: Study design, performed the research and managed the overall project, involved in entire study, analyzed the data, made the figures, and wrote the manuscript

FS: Performed the experiments, data analysis, made the figures, wrote the manuscript

DACB: performed the experiments and data analyses

KSL, NC, MSS, OR: provided tools, data analyses and discussions, funding generation and edited the manuscript

