## Supplementary material for "Marker-free imaging of α-Synuclein aggregates in a rat model of Parkinson’s disease using Raman microspectroscopy": Suppl. Figures and texts

**Suppl. Table 1: List of chemicals used for the study**

| Chemical | Company | Order Number |
|---|---|---|
| Bovine Serum Albumin | Sigma Aldrich (Darmstadt, GER) | A9576 |
| Cresyl Violett Acetate | Waldeck (Münster, GER) | 1A-400 |
| DAPI | Sigma Aldrich (Darmstadt, GER) | D9542 |
| Dulbecco's Phosphate-Buffered Saline (DPBS) | Gibco (Thermo Fisher Scientific) (Waltham, MA USA) | 14190-169 |
| Eosin G-Solution 1 % aqueous | Carl Roth (Karlsruhe, GER) | 3137.2 |
| Ethanol (EtOH) absolute | AppliChem (Darmstadt, GER) | A367812 |
| Gelatin from cold water fish skin | Sigma Aldrich (Darmstadt, GER) | G7041 |
| Goat Serum | Vector Laboratories (Burlingame, CA, USA) | S-1000-20 |
| Hematoxylin solution according to Mayer | Carl Roth (Karlsruhe, GER) | T865.1 |
| Isomount 2000 | VWR (Darmstadt, GER) | 05547535 |
| Isopropanol (2-propanol) | Honeywell (Morris Plains, NJ, USA) | 563935-1L |
| Paraformaldehyde (PFA) 4% | Sigma Aldrich (Darmstadt, GER) | 15812-7 |
| ProLong Gold Antifade Mountant | Invitrogen (Thermo Fisher Scientific) (Waltham, MA USA) | P36934 |
| Tissue-Tek O.C.T. | Sakura Finetek (Torrance, CA, USA) | 4583 |
| Triton X-100 | Sigma Aldrich (Darmstadt, GER) | T9284 |
| Tween 20 | Sigma Aldrich (Darmstadt, GER) | P1379 |

**Suppl. Table 2: List of antibodies used for the study**

| Name | Dilution | Company | Order Number |
|---|---|---|---|
| Primary Antibodies | | | |
| α-Synuclein anti-Mouse Rabbit IgG | 1:100 | Cell Signaling Technology (Cambridge, GB) | D37A6 |
| Collagen I A1 Mouse IgG1 | 1:800 | Novusbio (Centennial, CO, USA) | 113M4774 |
| Secondary Antibodies | | | |
| AlexaFluor® 488 goat anti-Rabbit IgG | | Invitrogen (Thermo Fisher Scientific) (Waltham, MA USA) | |

| | | | |
|---|---|---|---|
| | 1:250 | | A-11034 |
| AlexaFluor® 488 goat anti-Mouse IgG1 | 1:250 | Invitrogen (Thermo Fisher Scientific) (Waltham, MA USA) | A-21121 |
| AlexaFluor® 594 goat anti-Mouse IgG1 | 1:250 | Invitrogen (Thermo Fisher Scientific) (Waltham, MA USA) | A-21125 |
| AlexaFluor® 594 goat anti-Rabbit IgG | 1:250 | Invitrogen (Thermo Fisher Scientific) (Waltham, MA USA) | A-11037 |

**Suppl. Table 3: List of instruments used for data acquisition**

| Equipment | Company |
|---|---|
| OpticLab H850 Slide Scanner | plustek (Taipeh, TWN) |
| Axio Observer microscope Z1 | Zeiss (Oberkochen, GER) |
| Microscopy fridge | Liebherr (Bulle, CH) |
| Raman micro-spectroscope | WITec (Ulm, GER) |
| Laser scanning microscope | Zeiss (Oberkochen, GER) |
| Microm HM 560 Cryostat | Thermo Fisher Scientific (Waltham, MA, USA) |
| Microscope | Zeiss (Oberkochen, GER) |
| Suction device | neoLab (Heidelberg, GER) |

**Suppl. Table 4: List of software used for data analysis**

| Software | Developer | Application |
|---|---|---|
| Control Five | WITec (Ulm, GER) | Raman data acquisition |
| Project Five 5.2 | WITec (Ulm, GER) | Raman data processing |
| Zeiss (Zen) | Zeiss (Oberkochen, GER) | Microscope imaging program |
| ZEN 3.0 (blue edition) | Zeiss (Oberkochen, GER) | Microscope image processing |
| Microsoft Excel | Microsoft Corporation (Redmond, WA, USA) | Data analysis and processing |
| Microsoft Word | Microsoft Corporation (Redmond, WA, USA) | Text processing |
| The Unscrambler X 10.5 | CAMO Software (Oslo, NOR) | data analysis |
| GraphPad Prism 6 | GraphPad Software (San Diego, CA, USA) | Data analysis and processing |
| TeamViewer 14 | TeamViewer (Göppingen, GER) | Establishing connection from distance |

**Supp. Table 5: Identification of the Raman spectra**

The assigned spectra were listed along with their literature source

| SN | Wave number ($cm^{-1}$) | Assignment | References |
|---|---|---|---|

| | | | |
|---|---|---|---|
| 1 | 759 | Tryptophan | (1, 2) |
| 2 | 783 | DNA | (1, 3) |
| 3 | 790 | O-P-O stretching DNA | (1, 4) |
| 4 | 804 | Uracil-based ring breathing mode | (1) |
| 5 | 827 | Proline | (1) |
| 6 | 855 | Tyrosine | (1) |
| 7 | 860 | Phosphate group | (1) |
| 8 | 879 | Hydroxyproline | (1) |
| 9 | 938 | Proline | (1) |
| 10 | 957 | Cholesterol | (1) |
| 11 | 990 | C-C stretching | (1) |
| 12 | 1001 | Phenylalanine | (1) |
| 13 | 1032 | C-C stretching | (1) |
| 14 | 1053 | C-O stretching | (1, 5) |
| 15 | 1063 | C-C skeletal stretching | (1) |
| 16 | 1087-1099 | Phosphodioxy group (DNA backbone) | (1) |
| 17 | 1129 | C-C acyl backbone in lipid | (1) |
| 18 | 1156 | C-C/C-N stretching (proteins) | (1) |
| 19 | 1174 | Tyrosine | (1) |
| 20 | 1206 | Hydroxyproline, Tyrosine | (1) |
| 21 | 1248-1263 | Amide III, β-sheet | (6-10) |
| 22 | 1274 | Amide III, α-helix | (11) |
| 23 | 1280 | Amide III, α-helix | (1, 2) |
| 24 | 1290-1317 | Amide III, α-helix | (1, 12) |
| 25 | 1336-1342 | Amide III, α-helix | (10, 13) |
| 26 | 1378 | T, A, G (ring breathing modes of the DNA/RNA bases) | (1) |
| 27 | 1402 | C=O stretching | (1) |
| 28 | 1416 | C=C stretching in quinoid ring | (1) |
| 29 | 1440-1451 | $CH_2$ and $CH_3$ deformation | (1) |
| 30 | 1488 | Nucleic acid purine bases | (1) |
| 31 | 1570-1579 | Nucleic acids | (1) |
| 32 | 1586 | C=C olefinic stretching | (1) |
| 33 | 1592 | C=N and C=C stretching in quinoid ring | (1) |
| 34 | 1606 | C=C phenylalanine stretching | (1) |
| 35 | 1632-1637 | β-sheet | (8, 9, 14) |
| 36 | 1650-1658 | Amide I, α-helix | (15, 16) |
| 37 | 1664-1669 | Amide I, β-sheet | (10, 14) |
| 38 | 1670-1680 | Amide I, extended β-strand and polyproline II (PPII) structure | (1, 14, 17) |
| 39 | 1685 | Amide I, β-turn | (1, 14, 18) |
| 40 | 1706 | C=O stretching | (1) |

**Suppl. Figure legends**

**Suppl. Figure 1.** Identification of the area of interest in the brain sample. (a) Representative Nissl-stained brain section with the black square identifying the olfactory bulb area, also observed in the laser scanning microscope LSM. (b) Alpha-synuclein stained (D37A6) brain section with olfactory bulb. The area was visualized with the LSM. (c) Representative example of single spectrum scans in brain with antibody staining (D37A6) as shown in b. Single spectra measurements were depicted as crosses and large are scans as squares in the stained regions. Single spectra were used in spectral analysis and PCA analysis. The Raman spectra were identified based on different sources in literature. The exemplary spectrum was taken from WT 12M. (d) Identification of the Area of Interest in the colon sample WT 4M. H&E-stained colon section with the black square identifying the area in the 63x magnification. Magnification of 10x, pictured through the Raman microspectroscopy with the black square identifying the area in the 63x magnification. Magnification of 63x, pictured through the Raman microspectroscopy with the red square identifying the measured area with a large area scan. The layers of the colon tissue were identified as ME: Muscularis Externa, SM: Submucosa, MM: Muscularis Mucosa and E: Epithelial cells. Representative example of single spectrum scans with antibody staining (D37A6) in colon (right hand side image). Single spectra measurements were depicted as crosses in the most stained regions and used in spectral analysis and PCA analysis.

**Suppl. Figure 2.** Spectral comparison in the olfactory brain regions. (a) The WT 12M and TG 12M samples were compared through their mean and standard deviation. The intensities of the spectra were statistically analyzed through a t-test to detect any statistical changes. Differences were identified in the 759 cm-1 (p=0.04), 830 cm-1 (p=0.02), 877 cm-1 (p=0.04), 1268 cm-1 (p=0.008) and 1298 cm-1 (p=0.01) peaks.

**Suppl. Figure 3.** Comparisons of PCA for genotyping in the rat brain region. (a) Comparison of 12M WT and TG samples through PCA with scores and loadings in brain. The loadings of PC-6 were visualized.

**Suppl. Figure 4.** Comparisons of spectra and PCA for genotyping and ageing in the rat colon region. (a) FWHM spectra for 12M WT and TG colon samples. (b) Ageing comparison (4M vs 12M TG) of Raman spectra for normalized average height. (c) Comparison of 4M WT and TG samples through PCA with scores and loadings in colon. The 4M WT and TG samples were compared with PCA at PC-4 and PC-2. PC-4 was significant (p=0.001). The loadings of PC-4 were visualized. (d) Comparison of the amide III and amide I 4M WT and TG samples through PCA with scores and loadings in colon. The 4M WT and TG amide III samples were compared with PCA at PC-1 and PC-4. Both PC-1 (p=0.003) and PC-4 (p=0.01) were significant. The loadings of amide III, PC-1 and PC-4 were visualized. The 4M WT and TG amide I samples were compared with PCA at PC-2 and PC-7. Both PC-2 (p=0.03) and PC-7 (p=0.003) were significant. The loadings of amide I, PC-2 and PC-7 were visualized.

**Suppl. Figure 5.** Comparisons of PCA for genotyping (12M WT and TG) and ageing (3M-12M TG) in the rat colon region. (a) Comparison of the amide III 12M TG and WT samples through PCA with scores and loadings in colon. The 12M TG and WT amide III samples were compared with PCA at PC-5 and PC-6. PC-6 was significant (p=0.03). The loadings of amide III, PC-6 were visualized. (b) The 4M and 12M TG samples were compared with PCA at PC-2 and PC-4. Both PC-2 (p=0.008) and PC-4 (p=0.0004) were significant. Furthermore, 4M and 12M TG samples were compared with PCA at PC-3 and PC-4. Both PC-3 (p=0.03) and PC-4 (p=0.0004) were significant. The loadings of PC-2, PC-3 and PC-4 were visualized. (c) Comparison of the amide III and amide I 4M and 12M samples through PCA with scores and loadings in colon. 4M and 12M TG amide III samples were compared with PCA at PC-1 and PC-5. Both PC-1 (p=0.0003) and PC-5 (p=0.0007) were significant. 4M and 12M TG amide III samples were compared with PCA at PC-1 and PC-2. Both PC-1 (p=0.0003) and PC-2 (p=0.04) were significant. The loadings of amide III, PC-1, PC-2 and PC-5 were visualized. Furthermore, 4M and 12M TG amide I samples were compared with PCA at PC-2 and PC-3. Both PC-2 (p=0.00003) and PC-3 (p=0.03) were significant. Additionally, 4M and 12M amide I TG samples were compared with PCA at PC-2 and PC-5. Both PC-2 (p=0.00003) and PC-5 (p=0.004) were significant. The loadings of amide I, PC-2, PC-3 and PC-5 were visualized.

**Suppl. Figure 6.** TCA components in colon viewed in different groups. The components displaying the same results were put together and averaged in their groups. The five components calculated in all animal samples were collagen fibers, lipids, unknown component, cells and muscle fibers. The separated groups were WT 4M (red), TG 4M (blue), WT 12M (green), TG 12M (brown). Statistical ANOVA analysis was performed for intensity and FWHM and the significant peaks were presented in black squares. (a) Differences in collagen fibers were observed for intensity at 1245 cm-1 between WT 4M and 12M (p<0.0001) and WT and TG 12M (p<0.0001), and for FWHM at 759 cm-1 between TG 4M vs. 12M (p=0.02), at 1245 cm-1 between WT 4M vs. 12M (p=0.01) and WT vs. TG 12M (p=0.004), and at 1668 cm-1 between WT 4M vs. 12M (p=0.002). (b) Differences in lipids were observed for intensity at 1257 cm-1 between WT 4M vs. 12M (p<0.0001) and WT vs. TG 12M (p<0.0001), and for FWHM at 851 cm-1 between WT vs. TG 12M (p=0.03). (c) Differences in the unknown component were observed for intensity at 1004 cm-1 between TG 4M vs. 12M (p=0.02), and for FWHM at 1313 cm-1 between WT vs. TG 4M (p=0.0008). (d) Differences in cells were observed for intensity at 786 cm-1 between TG vs. WT 4M (p=0.04) and at 1665 cm-1 between TG vs. WT 4M (p=0.0006), and for FWHM at 1665 cm-1 between TG 4M vs. 12M (p=0.01). (e) Differences in muscle fibers were observed for intensity at 1004 cm-1 between TG vs. WT 4M (p<0.0001) and TG 4M vs. 12M (p=0.0002) and at 1653 cm-1 between TG vs. WT 12M (p=0.03). (f) Averaged intensity per pixel of muscle fibers statistically compared through t-test. The four groups were statistically compared through t-test. Differences were observed in muscle fibers between TG 12M vs. TG 4M (p=0.04) and TG 4M vs. WT 4M (p=0.057).

5. R. L. Frost, W. Martens, J. T. Kloprogge and Z. Ding: Raman spectroscopy of selected lead minerals of environmental significance. Spectrochimica Acta Part A:

Molecular and Biomolecular Spectroscopy, 59(12), 2705-2711 (2003) doi:10.1016/s1386-1425(03)00054-4
6. I. H. Chou, M. Benford, H. T. Beier, G. L. Cote, M. Wang, N. Jing, J. Kameoka and T. A. Good: Nanofluidic biosensing for beta-amyloid detection using surface enhanced Raman spectroscopy. Nano Lett, 8(6), 1729-35 (2008) doi:10.1021/nl0808132
7. S. M. A, R. Crockett, G. Kearn, I. Cherny, E. Gazit, C. S. L and P. J. S: Mechanically functional amyloid fibrils in the adhesive of a marine invertebrate as revealed by Raman spectroscopy and atomic force microscopy. Arch Histol Cytol, 72(4-5), 199-207 (2009) doi:10.1679/aohc.72.199
8. A. L. Fink: Protein aggregation: folding aggregates, inclusion bodies and amyloid. Folding and Design, 3(1), R9-R23 (1998) doi:10.1016/s1359-0278(98)00002-9
9. H. J. Lee, C. Choi and S. J. Lee: Membrane-bound alpha-synuclein has a high aggregation propensity and the ability to seed the aggregation of the cytosolic form. J Biol Chem, 277(1), 671-8 (2002) doi:10.1074/jbc.M107045200
10. C. Mensch, A. Konijnenberg, R. Van Elzen, A.-M. Lambeir, F. Sobott and C. Johannessen: Raman optical activity of humanα-synuclein in intrinsically disordered, micelle-boundα-helical, molten globule and oligomericβ-sheet state. Journal of Raman Spectroscopy, 48(7), 910-918 (2017) doi:10.1002/jrs.5149
11. M. Harris, K. Cilwa, E. A. Elster, B. K. Potter, J. A. Forsberg and N. J. Crane: Pilot study for detection of early changes in tissue associated with heterotopic ossification: moving toward clinical use of Raman spectroscopy. Connect Tissue Res, 56(2), 144-52 (2015) doi:10.3109/03008207.2015.1013190
12. E. W. Blanch, L. A. Morozova-Roche, L. Hecht, W. Noppe and L. D. Barron: Raman optical activity characterization of native and molten globule states of equine lysozyme: comparison with hen lysozyme and bovine alpha-lactalbumin. Biopolymers, 57(4), 235-48 (2000) doi:10.1002/1097-0282(2000)57:4<235::AID-BIP5>3.0.CO;2-H
13. E. W. Blanch, L. Hecht and L. D. Barron: Vibrational Raman optical activity of proteins, nucleic acids, and viruses. Methods, 29(2), 196-209 (2003) doi:10.1016/s1046-2023(02)00310-9
14. J. T. Pelton and L. R. McLean: Spectroscopic methods for analysis of protein secondary structure. Anal Biochem, 277(2), 167-76 (2000) doi:10.1006/abio.1999.4320
15. N. C. Maiti, M. M. Apetri, M. G. Zagorski, P. R. Carey and V. E. Anderson: Raman spectroscopic characterization of secondary structure in natively unfolded proteins: alpha-synuclein. J Am Chem Soc, 126(8), 2399-408 (2004) doi:10.1021/ja0356176
16. M. M. Apetri, N. C. Maiti, M. G. Zagorski, P. R. Carey and V. E. Anderson: Secondary structure of alpha-synuclein oligomers: characterization by raman and atomic force microscopy. J Mol Biol, 355(1), 63-71 (2006) doi:10.1016/j.jmb.2005.10.071
17. F. Zhu, J. Kapitan, G. E. Tranter, P. D. Pudney, N. W. Isaacs, L. Hecht and L. D. Barron: Residual structure in disordered peptides and unfolded proteins from multivariate analysis and ab initio simulation of Raman optical activity data. Proteins, 70(3), 823-33 (2008) doi:10.1002/prot.21593
18. T. Lefevre, M. E. Rousseau and M. Pezolet: Protein secondary structure and orientation in silk as revealed by Raman spectromicroscopy. Biophys J, 92(8), 2885-95 (2007) doi:10.1529/biophysj.106.100339

a

Brain section - Olfactory bulb

b

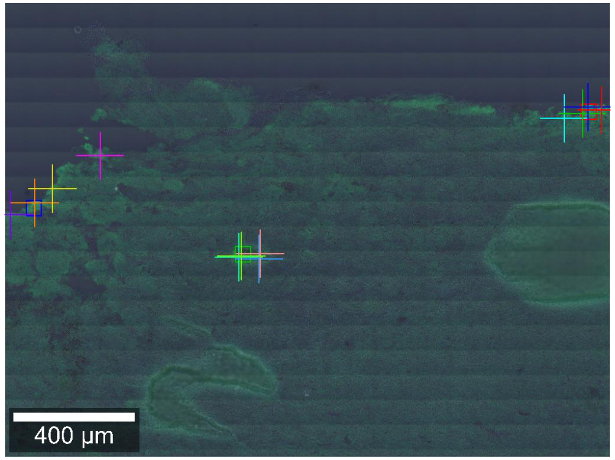


c

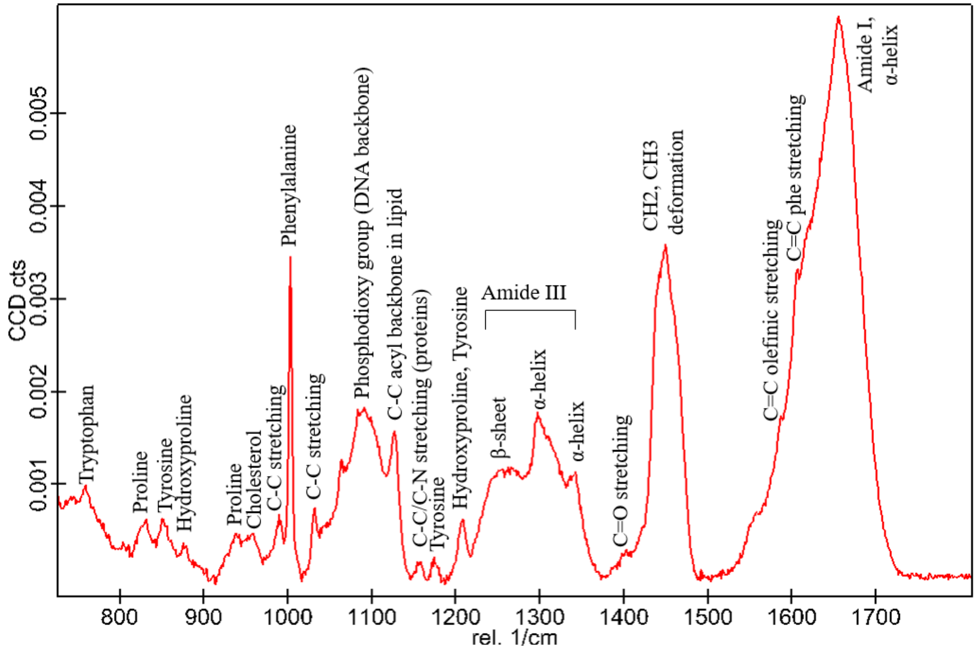


d

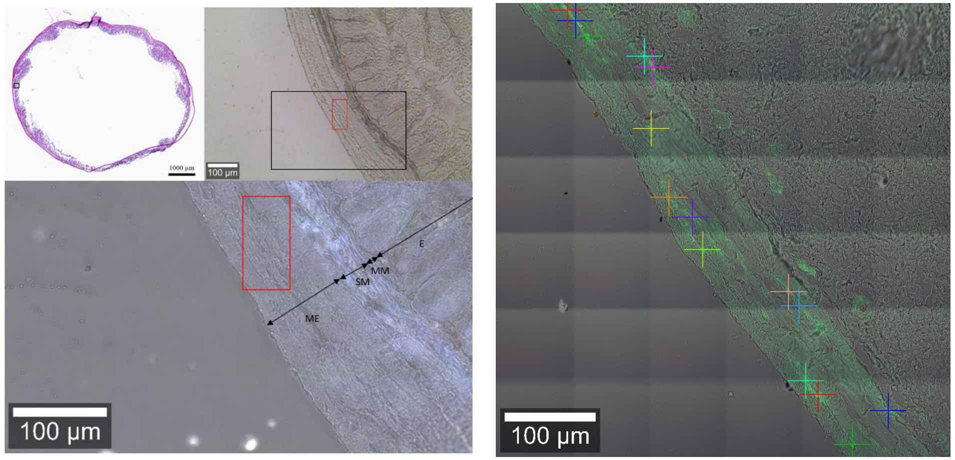


Suppl. Fig. 1

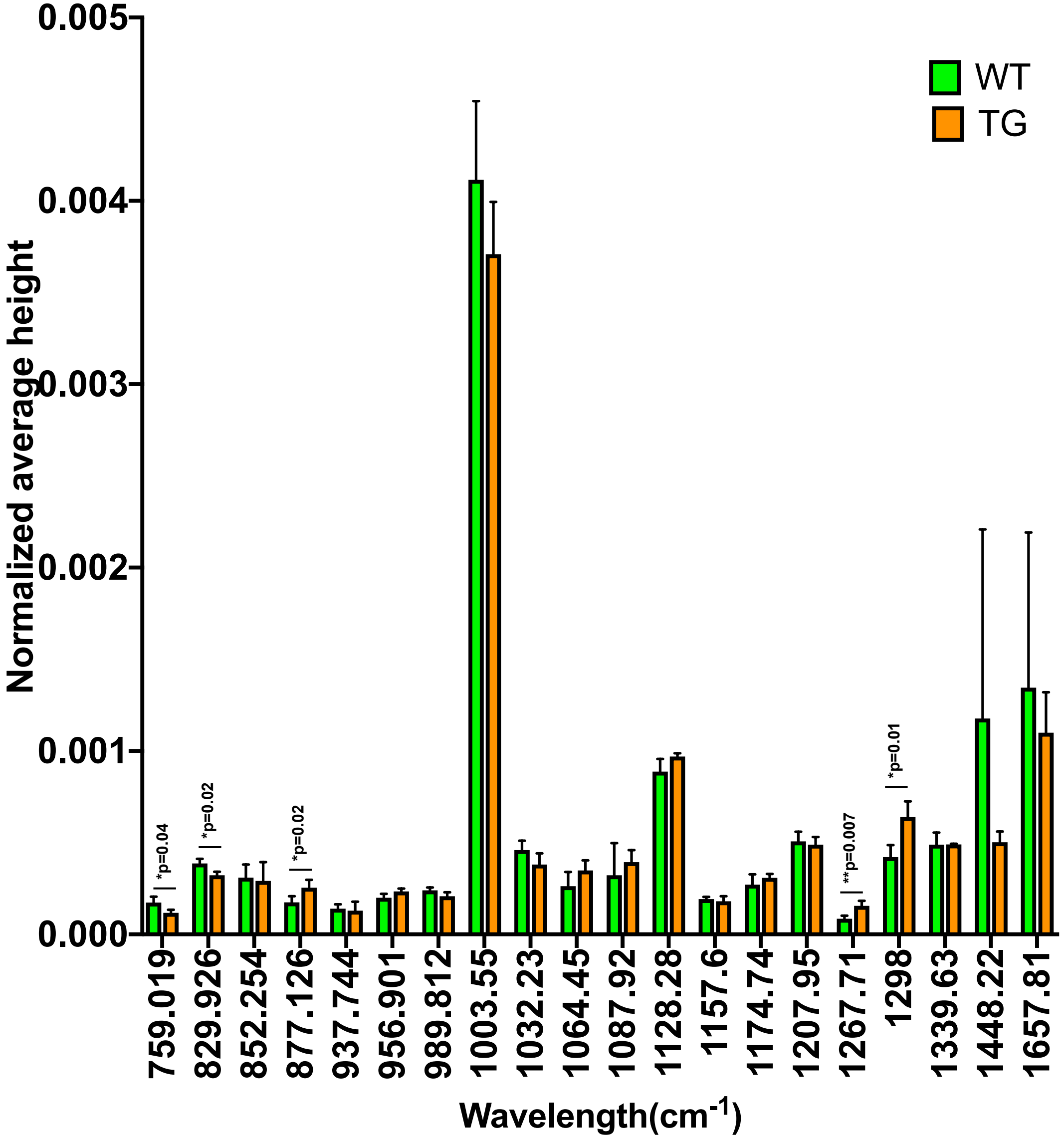
0.005
0.004
0.003
0.002
0.001
0.000
Normalized average height
WT
TG
*p=0.04
*p=0.02
*p=0.02
**p=0.007
*p=0.01
759.019
829.926
852.254
877.126
937.744
956.901
989.812
1003.55
1032.23
1064.45
1087.92
1128.28
1157.6
1174.74
1207.95
1267.71
1298
1339.63
1448.22
1657.81
Wavelength(cm-1)

Suppl. Fig. 2

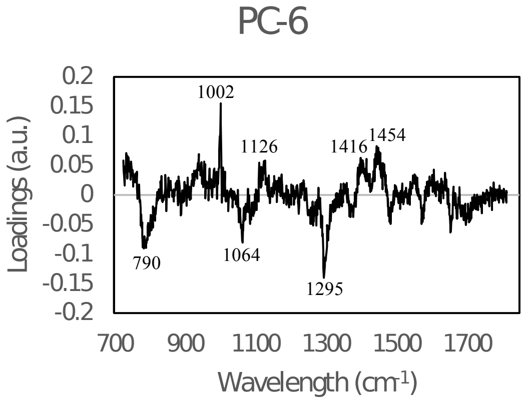
PC-6
Loadings (a.u.)
0.2
0.15
0.1
0.05
0
-0.05
-0.1
-0.15
-0.2
1002
1126
1416
1454
790
1064
1295
700
900
1100
1300
1500
1700
Wavelength (cm-1)

Suppl. Fig. 3

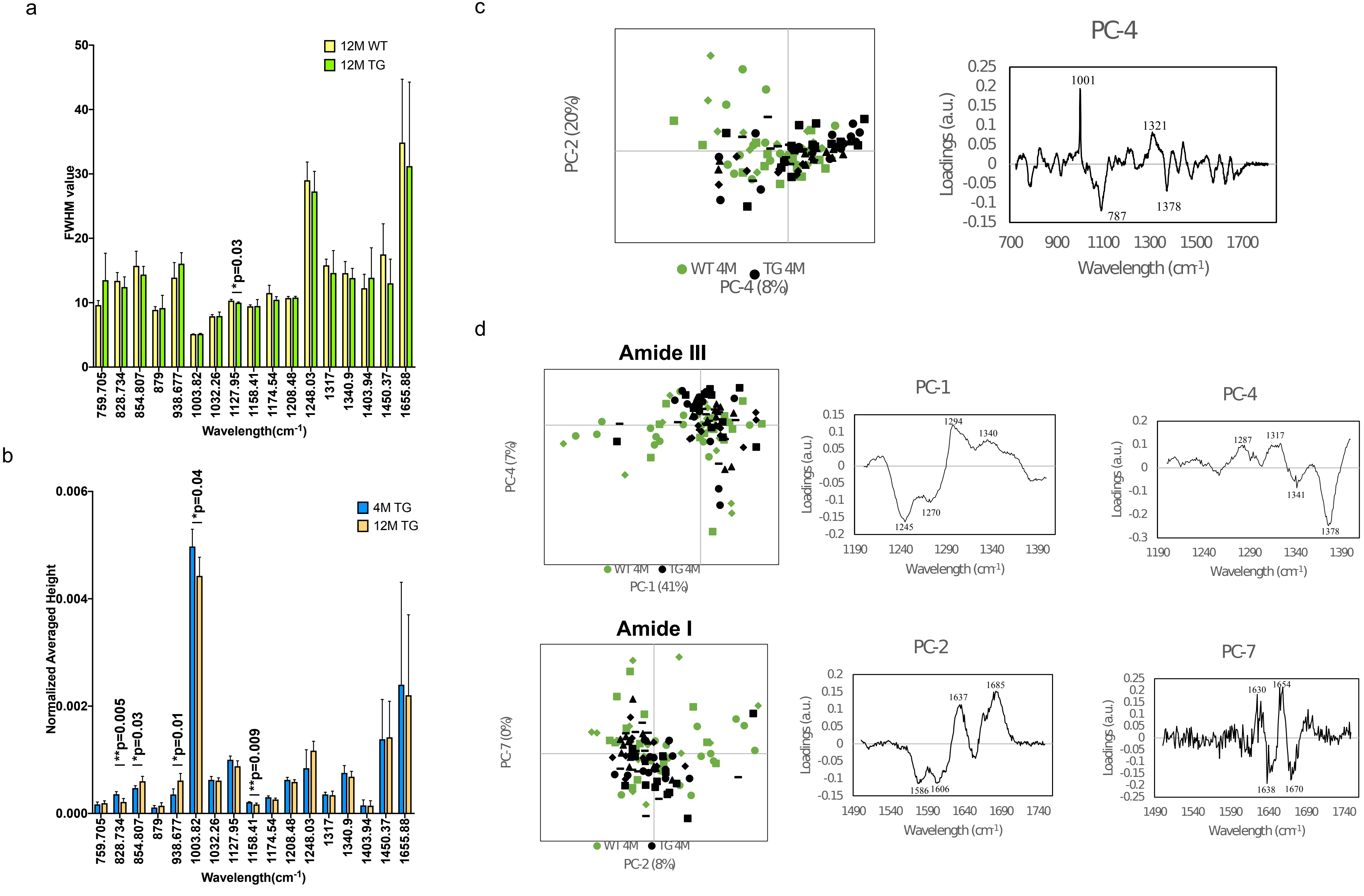

a
12M WT
12M TG
FWHM value
*p=0.03
Wavelength(cm-1)
b
4M TG
12M TG
Normalized Averaged Height
**p=0.005
*p=0.03
*p=0.01
*p=0.04
**p=0.009
Wavelength(cm-1)
c
PC-2 (20%)
WT 4M
TG 4M
PC-4 (8%)
PC-4
Loadings (a.u.)
Wavelength (cm-1)
1001
1321
787
1378
d
Amide III
PC-4 (7%)
PC-1 (41%)
PC-1
1294
1340
1245
1270
PC-4
1287
1317
1341
1378
Amide I
PC-7 (0%)
PC-2 (8%)
PC-2
1637
1685
1586
1606
PC-7
1630
1654
1638
1670


Suppl. Fig. 4

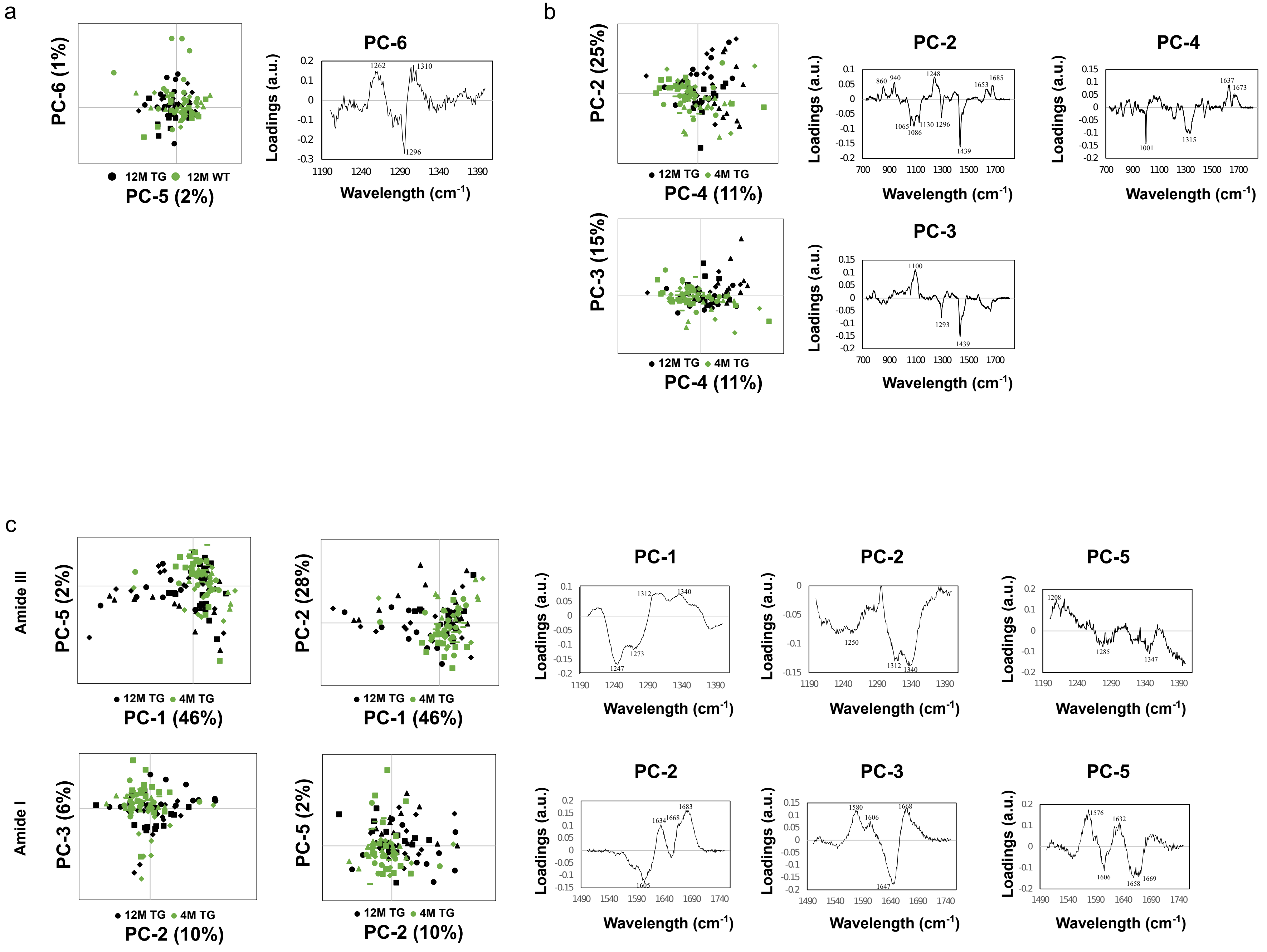

a
PC-6 (1%)
12M TG
12M WT
PC-5 (2%)
PC-6
Loadings (a.u.)
Wavelength (cm-1)
b
PC-2 (25%)
12M TG
4M TG
PC-4 (11%)
PC-2
PC-4
PC-3 (15%)
PC-3
Loadings (a.u.)
Wavelength (cm-1)
c
Amide III
PC-5 (2%)
PC-2 (28%)
12M TG
4M TG
PC-1 (46%)
PC-1
PC-2
PC-5
Amide I
PC-3 (6%)
PC-5 (2%)
PC-2 (10%)
PC-2
PC-3
PC-5
Loadings (a.u.)
Wavelength (cm-1)


Suppl. Fig. 5

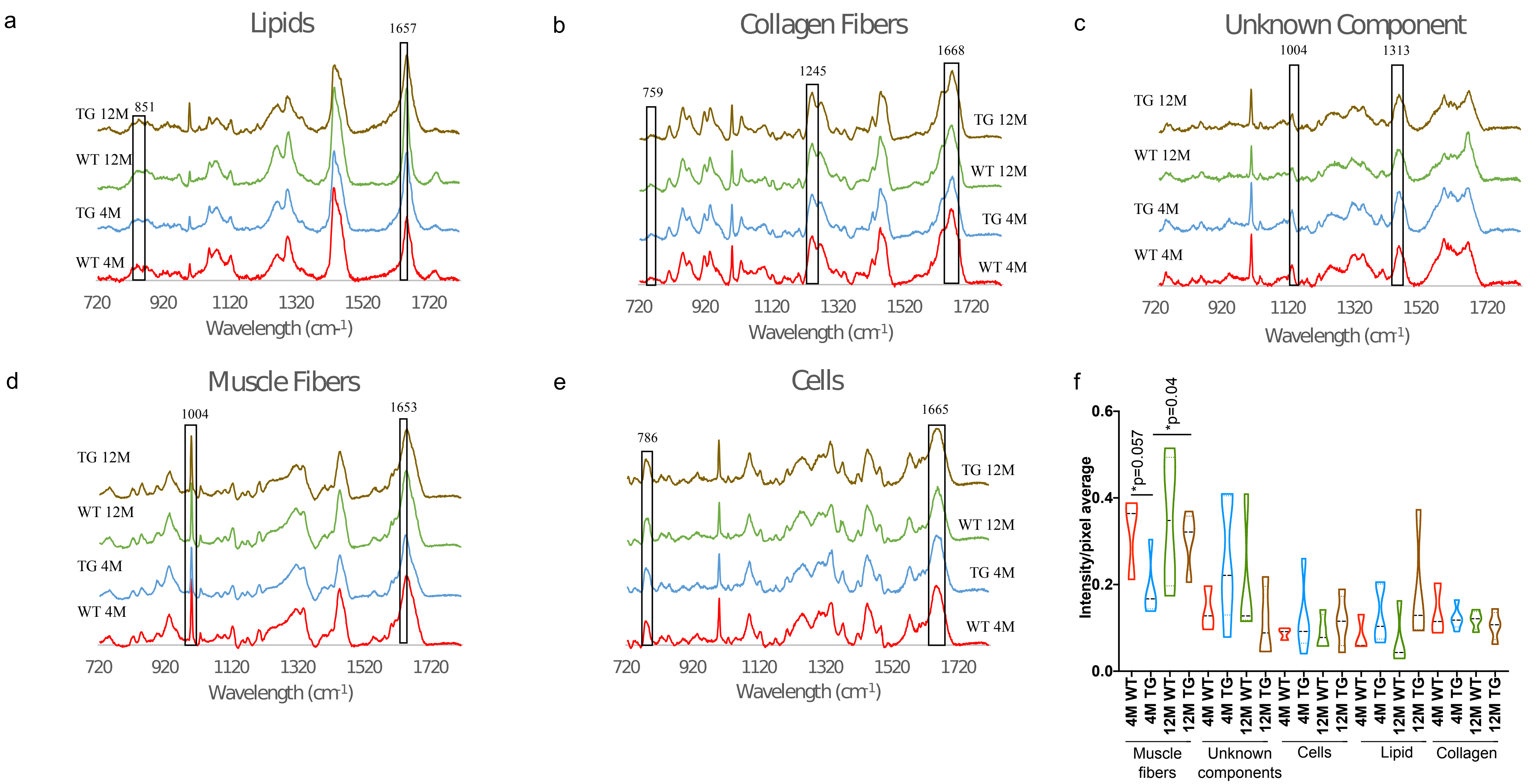
a
Lipids
1657
851
TG 12M
WT 12M
TG 4M
WT 4M
720
920
1120
1320
1520
1720
Wavelength (cm-1)
b
Collagen Fibers
759
1245
1668
TG 12M
WT 12M
TG 4M
WT 4M
720
920
1120
1320
1520
1720
Wavelength (cm-1)
c
Unknown Component
1004
1313
TG 12M
WT 12M
TG 4M
WT 4M
720
920
1120
1320
1520
1720
Wavelength (cm-1)
d
Muscle Fibers
1004
1653
TG 12M
WT 12M
TG 4M
WT 4M
720
920
1120
1320
1520
1720
Wavelength (cm-1)
e
Cells
786
1665
TG 12M
WT 12M
TG 4M
WT 4M
720
920
1120
1320
1520
1720
Wavelength (cm-1)
f
Intensity/pixel average
0.6
0.4
0.2
0.0
*p=0.057
*p=0.04
4M WT
4M TG
12M WT
12M TG
Muscle fibers
Unknown components
Cells
Lipid
Collagen

Suppl. Fig. 6
